# Dynamins combine mechano-constriction and membrane remodeling to enable two-step mitochondrial fission via a ‘snap-through’ instability

**DOI:** 10.1101/2024.08.19.608723

**Authors:** Haleh Alimohamadi, Elizabeth Wei-Chia Luo, Rena Yang, Shivam Gupta, Kelsey A Nolden, Taraknath Mandal, R. Blake Hill, Gerard C. L. Wong

## Abstract

Mitochondrial fission is controlled by dynamin proteins, the dysregulation of which is correlated with diverse diseases. Fission dynamins are GTP hydrolysis-driven mechanoenzymes that self-oligomerize into helical structures that constrict membrane to achieve fission, but details are not well understood. However, dynamins can also remodel membranes by inducing negative Gaussian curvature, the type of curvature required for completion of fission. Here, we examine how these drastically different mechanisms synergistically exert their influences on a membrane, via a mechanical model calibrated with small-angle X-ray scattering structural data. We find that free dynamin can trigger a “snap-through instability” that enforces a shape transition from an oligomer-confined cylindrical membrane to a drastically narrower catenoid-shaped neck within the spontaneous hemi-fission regime, in a manner that depends critically on the length of the confined tube. These results indicate how the combination of dynamin assembly, and paradoxically disassembly, can lead to diverse pathways to scission.

**Teaser:** Dynamin mechano-constriction by assembly and curvature-driven instability by free monomers synergistically drive mitochondrial fission.

## Introduction

The balance between the two antagonistic processes of mitochondrial fission and fusion is crucial in regulating the morphology and intracellular size distribution of mitochondria ^1–3^. Defects or disruptions of this balance impact metabolism and apoptosis, and have been correlated to developmental defects, cancer, cardiovascular diseases, and neurodegenerative diseases such as Parkinson, Alzheimers, Huntington, and amyotrophic lateral sclerosis (ALS) ^4–6^. The process of mitochondrial fission, in which a single mitochondrion divides into two separate daughter mitochondria, is facilitated by dynamin-related GTPase (Dnm1p) in yeast and its conserved human homologue (Drp1) ^1,7–10^. Dnm1p/Drp1 are GTPases from the dynamin-superfamily compromising of an N-terminal GTPase domain, a stalk or assembly domain, and an intrinsically disordered variable domain (IDVD) ^2,11^. In the current model, Dnm1p/Drp1 is generally acknowledged to oligomerize at the mitochondrial membrane surface, forming helical structures with an outer diameter of ∼50nm, with the GTPase domains facing the outside, and the IDVD domains on the inside, in contact with the mitochondrial membrane ^12–14^. Upon nucleotide hydrolysis, the Dnm1/Drp1 helical structures undergo a conformational change, locally constricting the mitochondrial membranes into a nanoscopic tube, and acts as a GTP hydrolysis-driven effector of fission ^12,15,16^. Interestingly, it has been experimentally observed that GTP hydrolysis also promotes disassembly of the oligomeric, confining helix ^17–19^. At present, the precise mechanism of mitochondrial fission is still a matter of debate considering constriction does not appear to lead deterministically to scission ^12^.

There are two main classes of extant models for Dnm1/Drp1 driven fission^12^. In ‘two-step models’, nucleotide-loaded dynamin assembles into helical structures, forming a scaffold around the mitochondrion to create a confined lipid tubule ^12,14,20^. GTP hydrolysis induces the subsequent disassembly of Dnm1/Drp1 oligomers ^17–19^ leading to the formation of hemi-fission intermediates and eventual scission ^12,14,20^. This composite process occurs via two time scales, a slow one for the confinement, and a fast one for the final scission. In contrast, for ‘constrictase models’, Dnm1/Drp1 acts like a molecular motor that enforces stepwise sliding of adjacent helical turns, leading to actively increasing degrees of constriction that itself leads to scission ^12^. Both models have their own explanatory power and limitations ^12^. To compound this already complex phenomenon, it has been noted that Dnm1 can induce negative Gaussian curvature (NGC) in membranes, the type of curvature necessary for scission ^21,22^.

Before we consider how mechanical confinement and curvature generation may or may not synergize with one another, it is critical to examine how these drastically different physical mechanisms exert their contrasting influences on a membrane. In recent years, theoretical and computational mechanical approaches informed by experimental data have recently become powerful tools for investigating the mechanisms underlying membrane remodeling and induced fission necks by dynamin protein machinery ^23–29^. For example, using coarse-grained molecular dynamic simulations, Pannuzzo et al. have proposed that the rotation of dynamin filaments around their longitudinal axis induces a local torque, which can trigger the topological transition of mitochondria into the hemi-fission state ^22^. In the continuum framework, previous studies have suggested that the localization of conical lipids in the pinching domain and the non-axisymmetric collar pressure induced by the helical arrangement of dynamin facilitate the membrane scission process and stabilize the highly constricted necks ^26,27,30^. Utilizing theoretical models and experimental measurements of ionic conductivity in a tubular membrane connected two parallel axisymmetric rings, Frolov et al. have demonstrated there is a critical tubule length at which the tubule transitions from a cylindrical nanotubule to a catenoid in less than a millisecond ^24^.

Here we provide a general framework to reconcile the two types of models and show how constrictase activity and hydrolysis driven disassembly synergistically contribute to fission.

Specifically, we use a mechanical model combined with small-angle X-ray scattering (SAXS) structural data to systematically investigate the role of NGC induced by dynamin and show its action on nanoscopic cylinders generated via various degrees of mechanical constriction by assembled dynamins. We find an unanticipated dynamin-induced “snap-through instability” that triggers a rapid shape transition from a cylindrical membrane tube to a highly constricted catenoid-shaped neck. This transition occurs within the range of the spontaneous hemi-fission regime. Importantly, whereas many studies have focused on the role played by the width of the dynamin oligomer confined membrane tube, we find that the length of this tube can be pivotally important: the occurrence of the dynamin-induced “snap-through” transition depends critically on the length of the confined membrane tube. We also show that the large anisotropic bending rigidity in conjunction with high membrane tension facilitates the mitochondrial fission process. These results suggest how dynamins’ mechanoenzyme activity to form nanoscopic tubes can synergize with their membrane remodeling activity, and thereby provide a point of contact to begin reconciliation of extant models. Finally, we apply our framework to compare the radius of fission necks induced by Dnm1 and Drp1 in various lipid compositions. Our results show that both Dnm1 and Drp1 can consistently induce narrow and robust necks in the hemi-fusion range, for lipid membranes with a high percentage of cardiolipin. This behavioral trend with cardiolipin from our data-calibrated mechanical model is consistent with previous experimental studies ^31–33^. Using coarse-grained molecular dynamic simulations, we also illustrate that the Dnm1/Drp1 complex can induce large anisotropic curvature in membranes with higher cardiolipin content. We think that our mechanical framework can be generalized to a broader range of protein-based fission machinery that can generate NGC on the membrane.

### Model development

#### Membrane mechanics

Our system comprises a tubular lipid membrane and rigid curvature-inducing dynamin-related GTPase proteins that are embedded in the membrane surface. We model the lipid bilayer as an elastic shell with negligible thickness compared to its bending ^34,35^. This allows us to describe the geometry of the membrane in terms of its mean (*H*) and deviatoric (*D*) curvatures, defined as

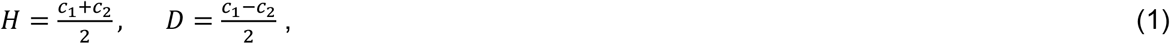

where *c*_1_ and *c*_2_ are the principal curvatures of the surface. For example, for an isotropic-shaped spherical bud with 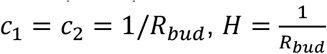 and *D* = 0. However, in the case of an anisotropic catenoid-shaped neck with *c*_1_ = −*c*_2_ = 1/*R*_*neck*_, *H* = 0 and 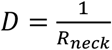.

We used the modified version of Helfrich energy that includes the deviatoric components to model the bending energy of the membrane, given as ^36–40^

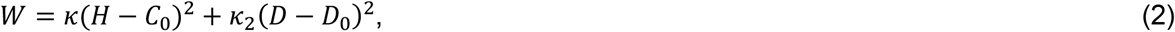

where *κ* is the membrane bending rigidity associated with isotropic curvature and *κ*_2_ is the membrane bending rigidity associated with anisotropic curvature. *C*_0_ is the induced spontaneous isotropic and *D*_0_ is the induced deviatoric curvatures by dynamin-related GTPase proteins. Assuming 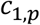 and 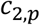 are the curvatures induced by dynamin in the two principal directions, 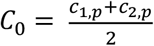 and 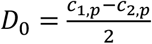. As an illustration, *C*_0_ captures the curvatures induced by proteins that form spherical coats ^41^, such as clathrin, while *D*_0_ represents the curvatures induced by proteins such as BAR domains and dynamins, which form tubular and neck-shaped structures. It should be mentioned that with no induced curvatures (*C*_0_ = 0 and *D*_0_ = 0), Eq. 2 reduces to the classical Helfrich energy with quadratic dependence on mean curvature and linear dependence on Gaussian curvature, where *κ*_2_ represents the magnitude of the Gaussian modulus (*k*_2_ = −*k*_*G*_).

Assuming the membrane is inextensible and the whole system of membrane and proteins is in mechanical equilibrium at all times, the normal force balance leads to the “shape equation” given as ^37,42,43^

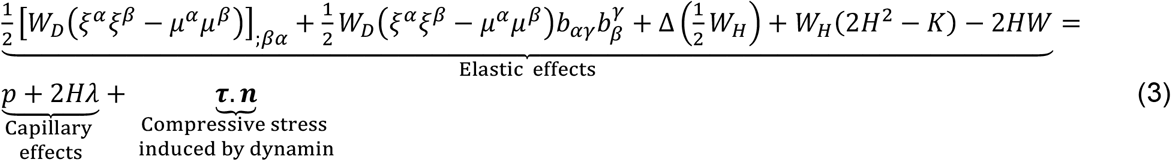

where *ξ*^*α*^ and *μ*^*α*^ are the projections of ***ξ*** and ***μ*** along the tangent vector on the surface. Here, ***ξ*** is a unit vector representing the orientation of protein, and ***μ*** is defined as ***μ*** = ***n*** × ***ξ*** where ***n*** is the unit surface normal ^40^ (Fig. 1B). *b*_*αγ*_ is the coefficients of the second fundamental form, 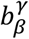 is the mixed components of the curvature, and (.)_;*α*_ is the covariant derivative. *W*_*D*_ and *W*_*H*_ are the partial derivatives of the energy density *W*, and *K* is the Gaussian curvature. Δ is the surface Laplacian, *λ* is the Lagrange multiplier for the area constraint interpreted to be the membrane tension ^44^, and *p* is the pressure difference across the membrane. ***τ*** is the compressive force density on the membrane induced by helical arrangements of dynamin proteins ^37^.

**Figure 1.**
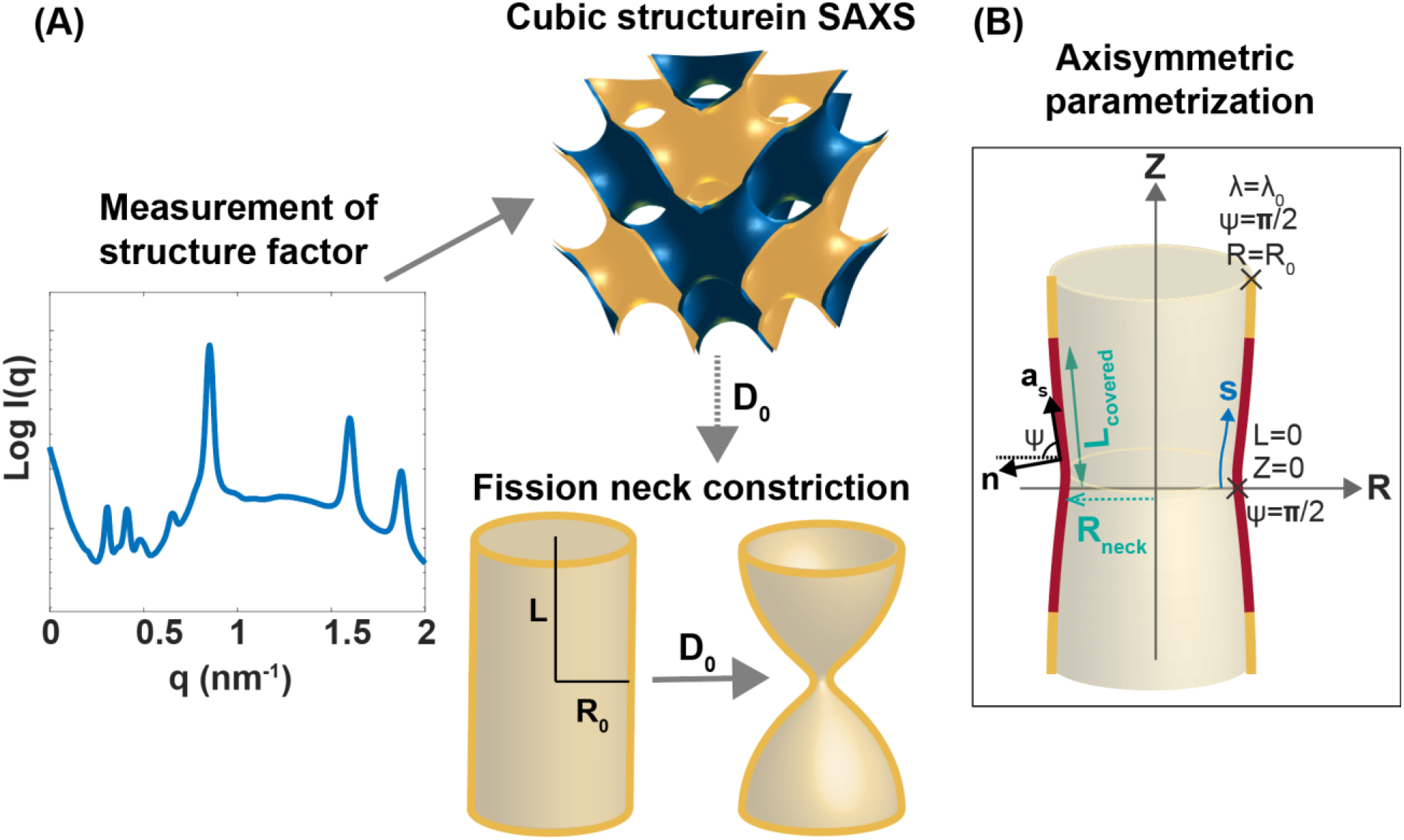
**(A)** Using induced membrane curvature deformations by dynamin-related proteins in SAXS spectra to estimate the radius of mitochondrial fission. The magnitude of generated anisotropic curvature by dynamin in cubic phases is used as an input for the mechanical model, constricting a tubular membrane with a radius of R_0_ = 20 nm and a half-length of L. **(B)** The surface parametrization of the tubular membrane with a radius of R_neck_ in axisymmetric coordinates and the prescribe boundary conditions. The yellow region represents the bare membrane, and the red region with a length of L_covered_ is the covered domain by dynamin. s is the arclength, ***n*** is the unit normal vector to the membrane surface, ***a***_***s***_ is the unit tangent vector in the direction of arclength, and ψ is the angle made by the tangent vector (***a***_***s***_) with respect to its radial plane.

A balance of forces tangent to the membrane yields the spatial variation of membrane tension given as ^37,42,43,45^

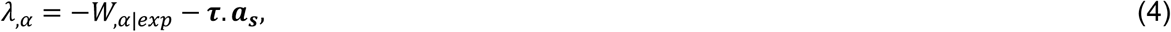

where (.)_*α*_ is the partial derivative, (.)_|*exp*_ denotes the explicit derivative with respect to coordinate, and ***a***_***s***_ is a tangent vector on the surface (Fig. 1B). The Supplementary Material provides a detailed derivation of the force balance’s governing equations and the procedure for non-dimensionalization.

The magnitude of induced deviatoric curvature (*D*_0_) in Eq. 2 can be estimated from the cubic structure formed by dynamin-related proteins in SAXS, given as ^22,46,47^

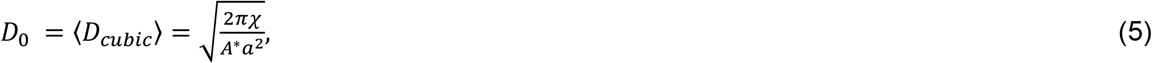

Where ⟨*D*_*cubic*_⟩ is the average membrane curvature deviator in a cubic phase, *a* is the lattice constant of the cubic phase, *χ* is the Euler characteristic, and *A*^*^ is the surface area per unit cell specific to each cubic phase ^48,49^.

#### Numerical implementation

For simplicity in the numerical calculations, we assumed that the tubular membrane is rotationally symmetric and also has a reflection symmetry with respect to the Z = 0 plane (Fig. 1B). In axisymmetric coordinates, the membrane shape equation (Eq. 3) and the tangential force balance equation (Eq. 4) reduce to a coupled system of first order differential equations (E. S12)^50^. Having the amount of induced deviatoric curvature from SAXS data (Eq. 5) as an input, we used “bvp4c” a boundary value problem solver in MATLAB, to solve this system of equations coupled with the prescribed boundary conditions shown in Fig. 1B (Eqs. S16 and S17) ^51,52^. In our simulations, we set *κ* = 10*kT* ^53^ and started from a tubular membrane with a radius of R_0_ = 20 nm^18,54,55^ and a half-length of L (Fig. 1B). In our study, we set *p* = 0 to focus only on the mechanism of dynamin-related proteins in governing the mitochondrial fission process. We also assumed that the magnitude of induced isotropic curvature by dynamin-related proteins is negligible compared to the magnitude of anisotropic curvature (*C*_0_ ≪ *D*_0_).

#### Induced anisotropic curvature by molecular motor dynamin can drive spontaneous mitochondrial fission via a snap-through transition

The dynamin family of proteins can constrict tubular membranes to form narrow necks. The size of these necks is a key geometric parameter that controls the dynamics of mitochondrial division, including the spontaneous fission process and the fission time from a few seconds to a couple of minutes ^12,56^. Here, we use our mechanical framework to understand how (*i*) the lattice constant of the induced cubic phases by dynamin, as observed in SAXS measurements ^21^, and (*ii*) the area of membrane tubule covered by dynamin (L_covered_) are correlated to the size of the mitochondrial constricted necks (Fig. 1C). To do that, we started from a tubular membrane with R_0_ = 20 nm, L/ R_0_ =2 and set *k*_2_/*k* = 1. In mechanical equilibrium and with no deviatoric curvature, the membrane tension required to maintain a tubular membrane (*λ*_*cylinder*_) depends on the bending rigidities and the tubule radius as 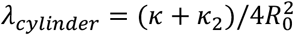(Eq. S19) ^37,57^. Here, we set the tension at the boundary as *λ*_0_ = *λ*_*cylinder*_.

In Fig 2A, we plotted the radius of the neck as a function of the protein coverage for three different *Pn3m* cubic lattice constants corresponding to different magnitudes of induced deviatoric curvature. We observed that for large lattice constants, e.g., *a* = 20 nm and *a* = 30 nm, the radius of the neck continuously decreases with an increase in the percentage of protein coverage (Fig. 2B). However, for a small lattice constant of *a* = 10 nm (large deviatoric curvature), the neck constriction with increasing protein coverage is associated with a snap-through transition from a wide neck (R_neck_ ∼ 8.5 nm) to a narrow neck (R_neck_ < 3 nm) in the limit of spontaneous hemi-fission ^58^ (Fig. 2A). The snap-through instability resembles the fast fission transition (less than 100 ms) induced by the GTP hydrolysis and disassembly of dynamin oligomers, as proposed in two-step dynamin’s fission model ^12^. Interesting, we observed that after the snap-through transition there is a minimum neck size of R_neck_ ∼ 2.6 nm, and then the size of the neck slightly increases with further increase in protein coverage (see the insert in Fig. 2A).

**Figure 2.**
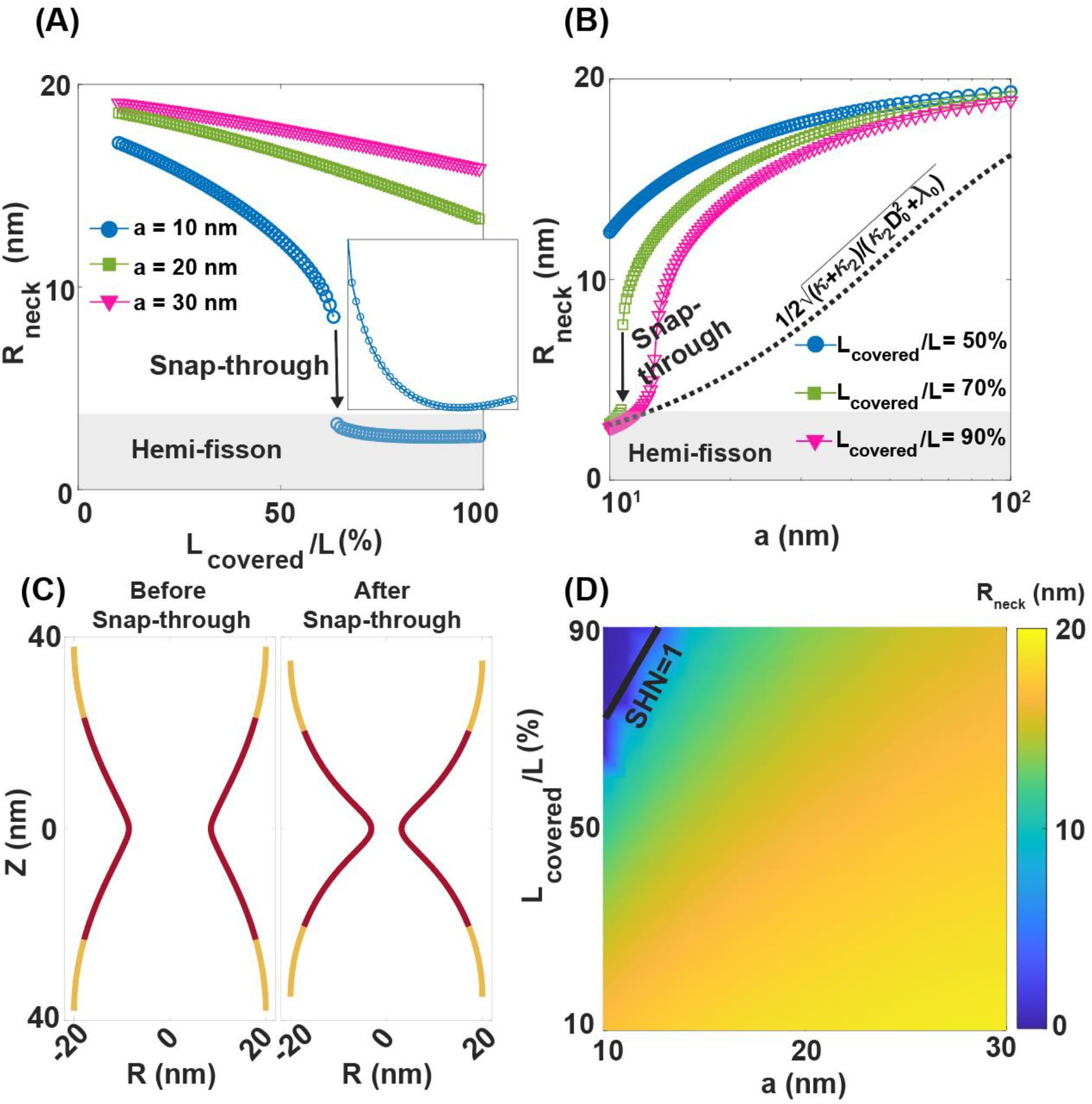
Estimating the radius of mitochondrial fission neck by dynamin-related proteins using the lattice constant of induced cubic structures in SAXS experiments. **(A)** The radius of the mitochondrial constricted neck as a function of protein coverage for three different cubic lattice constants. For a small lattice constant, there is a snap-through transition from a wide to narrow neck with increasing protein coverage. **(B)** The radius of the mitochondrial constricted neck as a function of the cubic lattice constant for three different protein coverages. As the lattice constant of the cubic phase decreases, the neck becomes narrower. The dotted black line represents to the analytical solution for the equilibrium radius of a tubular membrane in the presence of deviatoric curvature, given as 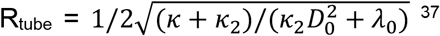^37^. **(C)** The morphology of the constricted neck before and after snap-through transition with a fixed *a* = 10 nm. **(D)** Phase diagram of constricted neck for a range of cubic lattice constants and protein coverages. The solid black line indicates SHN=1, above which R_neck_ < 3 nm within the spontaneous hemi-fission regime. *k*_2_/*k* = 1, *λ*_0_/*λ*_cylinder_ = 1, and L/R_0_ = 2.

We also plotted the radius of the mitochondrial neck as a function of the lattice constant of a *Pnm3* cubic phase for three different percentages of protein coverage. We observed the radius of the neck decreases with the decrease in the cubic lattice constant (larger induced deviatoric curvature in Eq. 4). This is in agreement with the analytical solution for the equilibrium radius of a tubular membrane in presence of spontaneous deviatoric curvature, given by 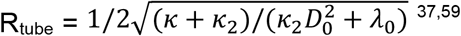 (dotted black line in Fig. 2B). Based on the length of the protein coverage, we identified three regimes of neck constriction: (*i*) For small protein coverage (L_covered_/L = 50%), the neck constriction is smooth, and a decrease in the cubic lattice constant from *a* = 100 nm to *a* =10 nm, results in a ∼36% reduction in the neck radius from R_neck_ ∼ 20 nm to R_neck_ ∼ 12.4 nm (Fig. 2B). (*ii*) For intermediate protein coverage (L_covered_/L = 70%), the neck constriction is associated with a snap-through transition and the radius of the neck decreases significantly, by ∼87%, reaching the spontaneous hemi-fission regime (R_neck_ < 3 nm) at small cubic lattice constants (Fig. 2B). (*iii*) For large protein coverage (L_covered_/L = 90%), the radius of neck decreases smoothly from R_neck_ ∼ 20 nm to R_neck_ ∼ 2.6 nm as the cubic lattice constant decreases from *a* = 100 nm to *a* =10 nm (Fig. 2B). The morphology of constricted membrane tubes before and after snap-through transition is shown in Fig. 2C. Before snap-through, the membrane tubule constricted to a wide neck, whereas a narrow, catenoid-shaped neck forms after the snap-through transition (Fig.2C).

In Fig. 2D, we plotted a phase diagram of the radius of the constricted neck for a range of cubic lattice constants and protein coverage. To estimate the range of spontaneous hemi-fission regime based on the magnitude of cubic lattice constant and the length of the protein coverage, we introduced a dimensionless quantity, the spontaneous hemi-fission number 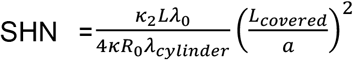. Based on our results, a SHN =1 is a threshold for the formation of narrow necks, while a SHN >1 corresponds to constricted necks within the spontaneous hemi-fission regime (above the solid black line in Fig. 2D). Overall, these results suggest that the anisotropic curvature induced by dynamin can remodel tubular membranes into narrow necks within the range of the spontaneous hemi-fission regime and our model predicts that this transition can be discontinuous, occurring through a snap-through instability.

#### Collar pressure induced by helical arrangements of the dynamin smoothly constricts mitochondrial membrane into narrow necks

It is well known that the helical assembly of the dynamin family of proteins can squeeze mitochondrial membranes to narrow necks, thereby facilitating the fission process ^16,60–62^. To compare the constriction mechanism of a tubular-shaped mitochondria by dynamin-induced anisotropic curvature (*D*_0_) with the collar pressure generated by rings of dynamin assembly (*τ*), we repeated the same simulation in Fig. 2 with no deviatoric curvature (*D*_0_ = 0). We assumed that the helical rings of dynamin generate a uniform compressive force density ***τ*** over a length of L_force_ on the tubular membrane (Fig. 3A). We found that the compressive force density induced by the helical orientation of dynamin can smoothly constrict the tubular membrane into a narrow neck and the degree of the constriction slightly depends on the length of the applied force density (Fig. 3A). This is consistent with the previous study by McDargh et al, that have suggested that the helical assembly of dynamin does not destabilize membrane during mitochondrial fission ^25^. Indeed, the smooth constriction is reminiscent of the slow pinching process (occurring over a period of seconds) that is mediated by dynamin scaffolding, as proposed in the two-step dynamin’s fission model ^12^. Based on our results, a compressive force density *τ* > 1.5 pN/nm^2^ is required to form narrow necks within the range of the spontaneous hemi-fission limit (Fig. 3A).

**Figure 3.**
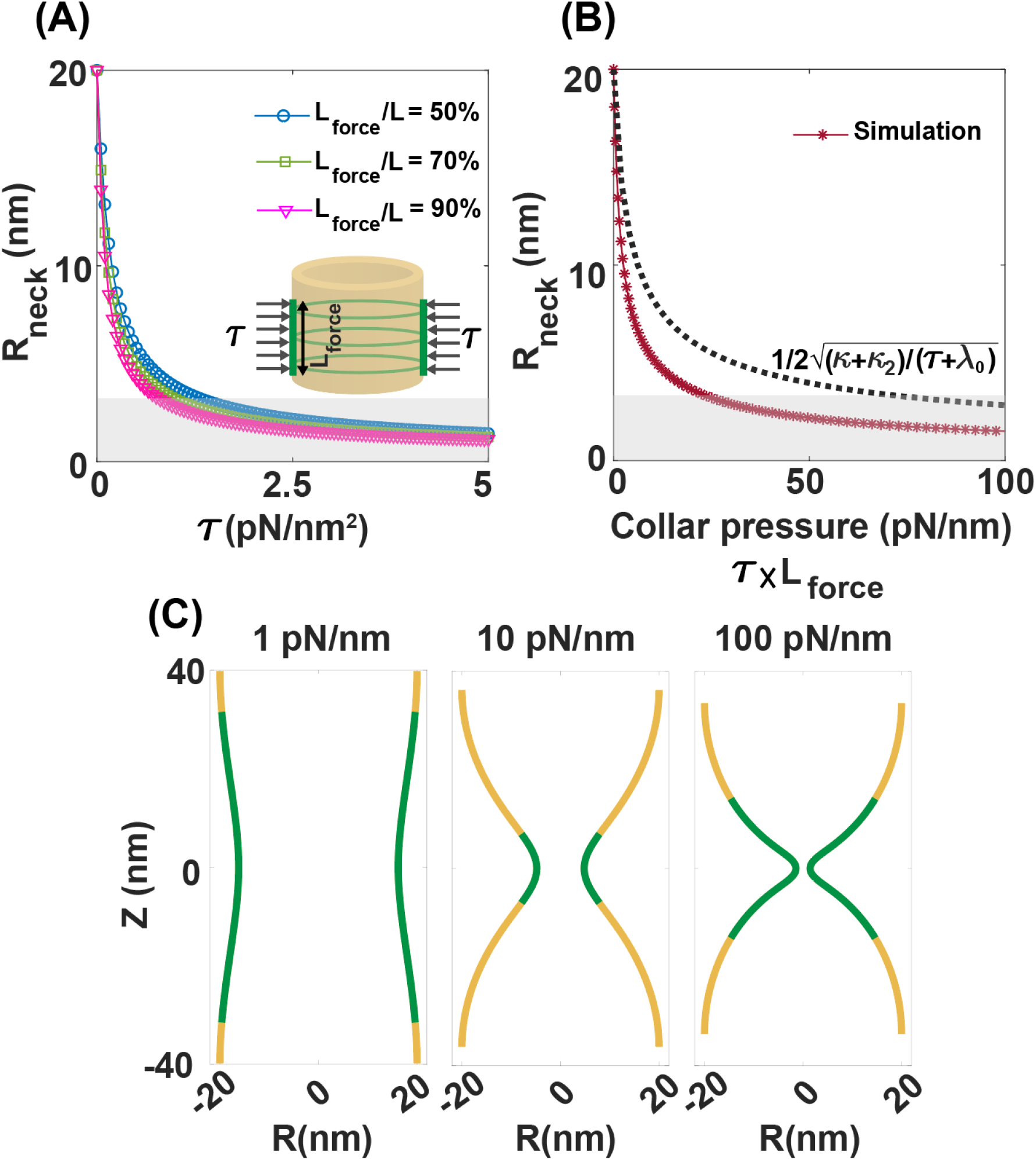
Continuous constriction of mitochondrial fission neck by compressive forces induced by dynamin helical rings (*D*_0_ = 0). **(A)** The radius of the mitochondrial constricted neck as a function of the compressive force density for three different lengths of the applied force. The insert shows the schematic of a tubular membrane with a uniform compressive force density (***τ***) applied over a length of L_force_ shown in green. **(B)** The radius of the mitochondrial constricted neck as a function of the collar pressure defined as *τ* × L_force_. The radius of the neck falls below the threshold for spontaneous hemi-fission for collar pressure > 25 pN/nm. The dotted black line corresponds to the analytical solution for the equilibrium radius of a tubular membrane under a uniform force density, given as 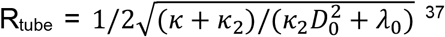. **(C)** The morphology of the constricted neck with three different magnitudes of the applied collar pressure. *k*_2_/*k* = 1, *λ*_0_/*λ*_cylinder_ = 1, and L/R_0_ = 2.

We combined the effects of force density and the length of the applied force by defining collar pressure as *τ* × L_force._ Based on our results, the radius of the neck decreases smoothly with an increase in the magnitude of the collar pressure, and a collar pressure > 25 pN/nm is needed to form narrow necks (R_neck_ < 3 nm) within the spontaneous hemi-fission regime (Fig. 3B). The analytical solution for the equilibrium radius of a tubular membrane under uniform force density 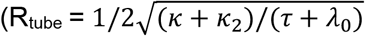, Eq. S21) is represented as a dotted black line (Fig. 3D). For small collar pressures, there is a good agreement between the analytical expression and numerical results (Fig. 3B). However, for large collar pressures, when catenoid-shaped necks form, the analytical solution deviates from the simulation results (Fig. 3B). In Fig. 3C, we demonstrate the morphologies of the constricted tubule under different magnitudes of applied collar pressure. The green line indicates the domain of applied compressive force (Fig. 3C). Using electron microscopy, a previous study by Stowell et al. suggested that a three-turn helix of dynamin can induce a compressive force of ∼330 pN ^63^. Based on our results, a collar pressure > 25 pN/nm is required for the formation of narrow necks in a tubular membrane (Fig. 3B). This implies that a dynamin helical structure, with more than one turn per ∼ 5 nm of membrane, is needed for successful mitochondrial fission. Overall, our findings in Figs. 2 and 3 can provide insight for the driving mechanisms and the relative time scales in each step of the two-step model of dynamin-catalyzed fission.

#### Longer mitochondrial membranes are easier to constrict by dynamin-induced anisotropic curvature

The length of tubular membranes formed by the dynamin helical structures can vary widely, ranging from nanometers to micrometers ^31,32^. Additionally, dynamin oligomerization and membrane constriction elongate tubular membranes confined by dynamin scaffolding ^63^. How does the length of the membrane tubule influence the mitochondrial fission rate? Using the mitochondrially targeted KikGR1 protein visualization technique, Cagalinec et al. have showed that in both in cortical and cerebellar granule neurons, the mitochondrial fission rate increased dramatically in longer mitochondria, while the fusion rate was almost independent of length ^64^. Berman et al have also proposed a model to relate the probability of a fission event in a mitochondria to its length ^65^. To explore the effect of mitochondria length on the constriction process induced by dynamin family of proteins, we repeated the simulations in Fig. 2 for a range of L/R_0_ (fixed R_0_ = 20 nm) from a short tubule (L/R_0_ = 0.5) to a long one (L/R_0_ = 4) (Fig. 4). We found that dynamin-induced anisotropic curvature (*D*_0_) can constrict longer tubules more effectively than shorter ones (Figs. 4B and 4C). For example, with a cubic lattice constant of *a* = 10 nm and L_covered_ = 50%, the induced curvature slightly reduces the neck size of a short tubule with L/R0 = 1 to R_neck_ ∼ 18 nm (Fig. 4B). However, for a long membrane tube with L/R0 = 4, the same curvature significantly constricts the tube into the spontaneous hemi-fission regime with R_neck_ < 3 nm (Fig. 4B). This could be attributed to the greater bending moment (force×length) induced by curvature-generating proteins in longer membrane tubules, which resembles the easier bending of a longer beam in classical mechanics.

**Figure 4.**
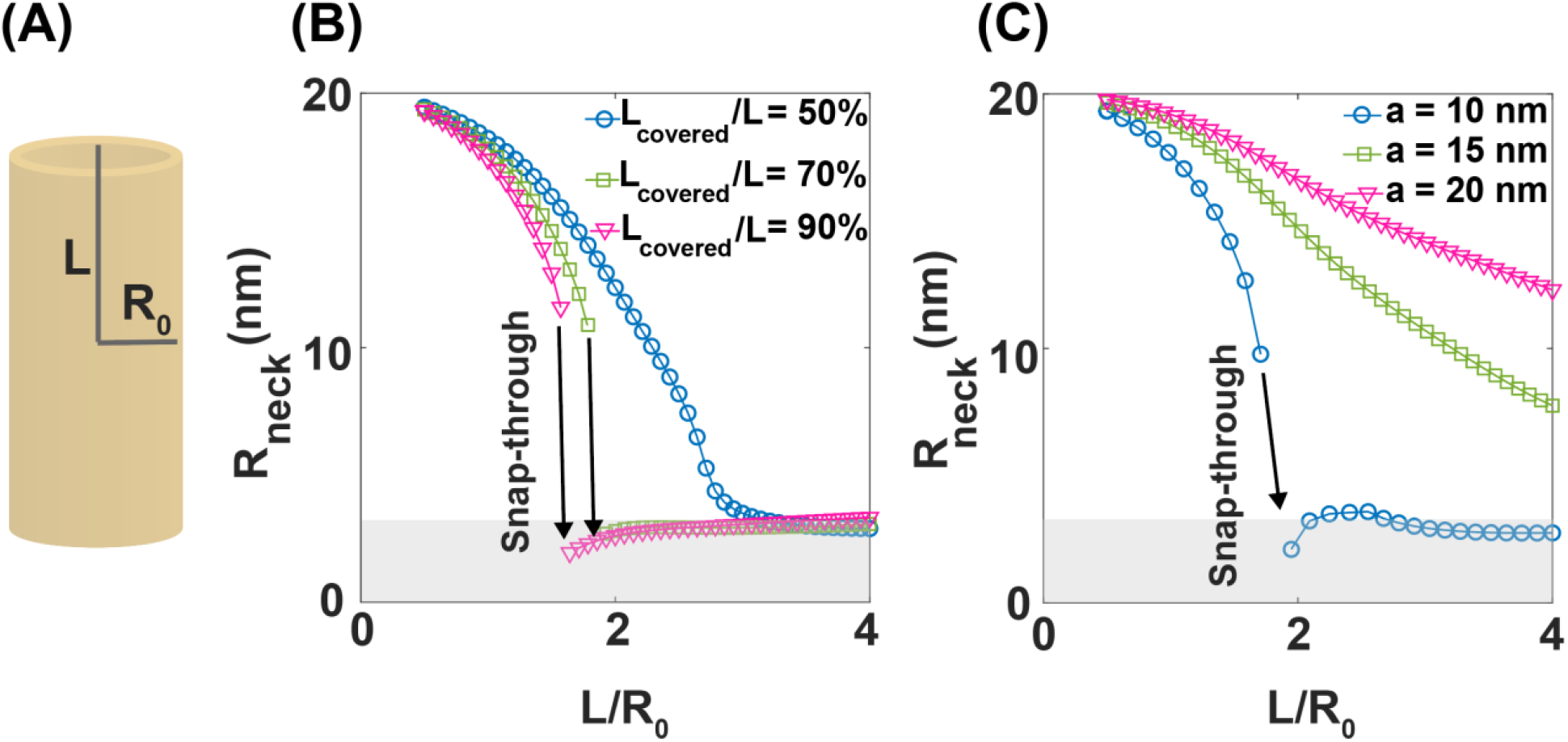
The length of mitochondrial affects the efficiency of dynamin-induced curvature constriction. **(A)** Schematic representation of a tubular membrane with a radius of R_0_ = 20 nm and a half-height of L. **(B)** The radius of the mitochondrial constricted neck as a function of the tubule height to the radius ratio (L/R_0_) for three different protein coverages (*a* = 10 nm). For large protein coverage (L_covered_/L = 70% and 90%), there is a discontinuous transition from a wide neck to the hemi-fission regime with an increase in the height of the membrane tubule. **(C)** The radius of the mitochondrial constricted neck as a function of L/R_0_ for three cubic lattice constants (L_covered_/L = 80%). For a small lattice constant (*a* = 10 nm), there is a discontinuous transition from a wide neck to the hemi-fission regime with an increase in the height of the membrane tubule. *k*_2_/*k* = 1, *λ*_0_/*λ*_cylinder_ = 1.

Interestingly, we observed that for large protein coverages and small cubic lattice constants, there is a snap-through transition in the radius of the constricted neck as L/R0 increases (Figs. 4B and 4C). Based on our results, the neck radius decreases smoothly when L/R_0_ increases from L/R_0_ = 0.5 to L/R_0_ < 2. Then, it abruptly transitions into the spontaneous hemi-fission regime at L/R_0_ ∼ 1.8, reaching the minimum neck size (Figs. 4B and 4C). Beyond this point, the neck size slightly increases with larger L/R_0_ ratios (Figs 4B and 4C). This snap-through instability resembles the spontaneous hemi-fission induced by GTP hydrolysis and sliding of adjacent turns of dynamin helices at a critical tubule length, as proposed in the constrictase dynamin’s fission model ^12^. Similarly, Frolov et al. have shown the shape bistability transition of cylindrical nanotubes into catenoidal microtubules is associated with an increase in the length of tubule lipid membranes ^24^. To investigate the net effect of tubule height on the efficiency of curvature-induced constriction, we kept the length of protein coverage constant (e.g., L_covered_ = 36 nm for all tubule heights which corresponds to 22.5% < L_covered_/L < 90%) and repeated the simulations (Fig. S1). Consistently, we found that the same amount of anisotropic curvature leads to the formation of a narrower constricted neck in longer tubules compared to shorter ones (Fig. S1). Thus, our mechanical framework suggests that longer mitochondrial membrane tubules formed by dynamin helicoidal assembly are susceptible to spontaneous hemi-fission triggered by dynamin-induced membrane remodeling machinery.

#### Interplay of membrane anisotropic bending rigidity and tension regulates the dynamic of mitochondrial constriction by dynamin-induced anisotropic curvature

The bending rigidity of the membrane describes its resistance to bending and it depends on the lipid composition of the membrane and the curvature generating proteins embedded within it ^66–69^. It has been demonstrated both theoretically ^70^ and experimentally ^68^ that a more negative Gaussian modulus (larger *k*_2_) in the protein-covered domain can facilitate membrane budding process and neck formation. However, measuring the Gaussian modulus of the lipid bilayer and protein composition at the neck domain remains challenging ^71^. Here, to understand how the magnitude of the anisotropic bending modulus relative to the isotropic bending rigidity (*k*_2_/*k*) affects the process of mitochondrial constriction by dynamin-induced deviatoric curvature, we conducted simulations across a wide range of *k*_2_/*k* and with different protein coverages (Fig. 5A, *a* = 10 nm).

**Figure 5.**
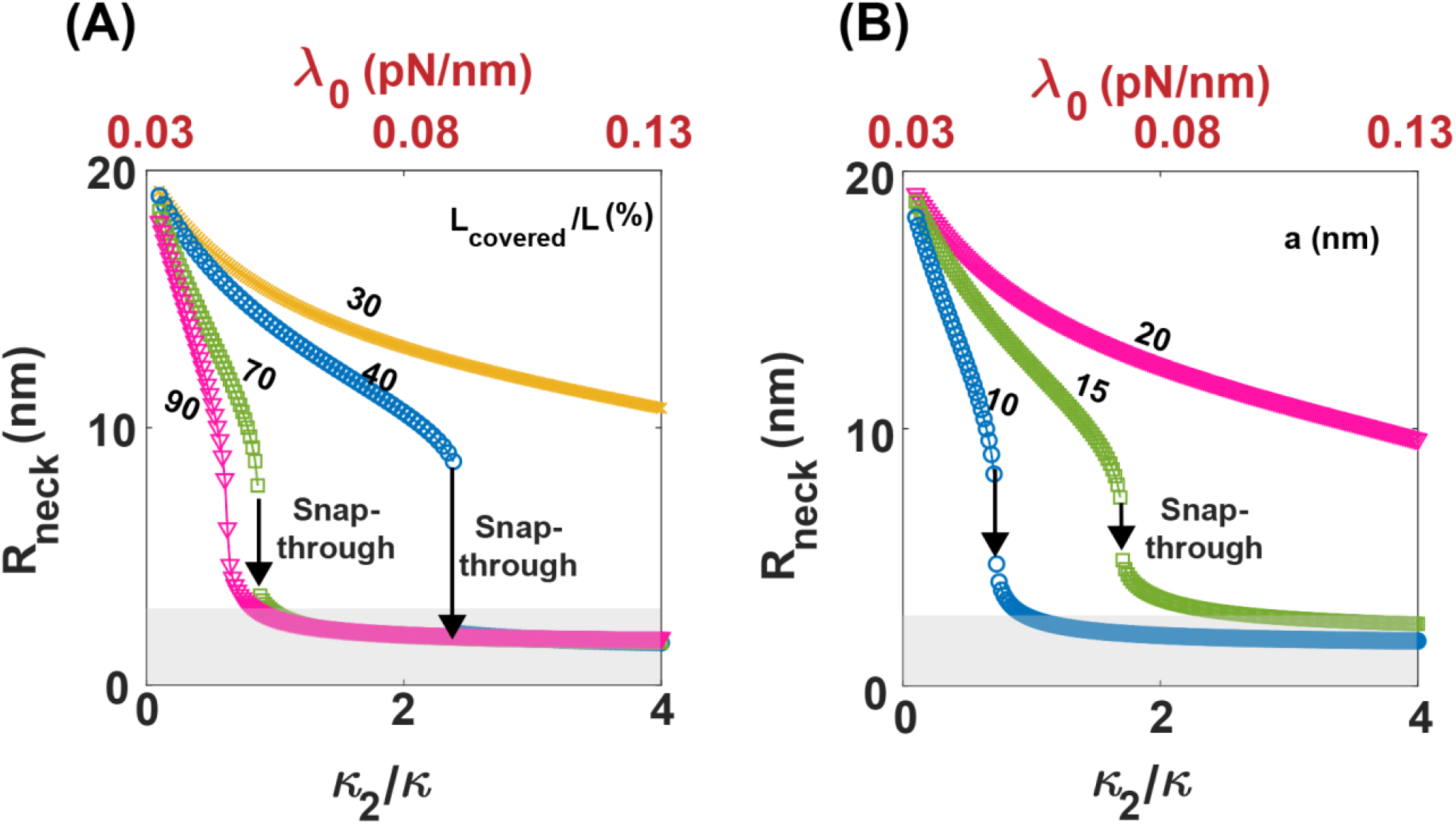
Membrane anisotropic bending rigidity coupled with tension controls the constriction of mitochondrial fission neck through dynamin-induced deviatoric curvature. **(A)** The radius of the constricted mitochondrial neck as a function of the anisotropic to isotropic bending moduli ratio (*k*_2_/*k*) for different protein coverages (*a* = 10 nm). We identified three distinct regimes. (*I*) For large protein coverage (L_covered_/L = 90%), the neck radius smoothly decreases into the hemi-fission regime as *k*_2_/*k* increases. (*II*) For intermediate protein coverages (40% < L_covered_/L <70%), there is a snap-through transition into the hemi-fission regime with increasing *k*_2_/*k*. (*III*) For small protein coverage (L_covered_/L = 30%), the neck radius slightly decreases as *k*_2_/*k* varies. The membrane tension at the boundary is set to be 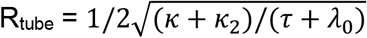 and the corresponding boundary tension for each *k*_2_/*k* value is marked at the top of the graph. **(B)** The radius of the constricted mitochondrial neck as a function of *k*_2_/*k* for three different cubic lattice constants (L_covered_ /L = 80%). For lattice constants of *a* = 10 and 15 nm, the transition into the spontaneous hemi-fission is associated with a snap-through instability. However, for a small lattice constant (*a* = 20 nm), the neck radius slightly decreases with an increase in *k*_2_/*k*. In all simulations, we set L/R_0_ = 2.

Our results showed that the necks become narrower with increasing membrane bending rigidity i.e. *k*_2_/*k*, which is consistent with previous studies ^68,70^. The comparison between the estimated neck radius from the analytical solution and the simulation results is shown in Fig. S2. Interestingly, we observed three different dynamics of neck constriction, depending on the percentage of protein coverage. (*i*) For large protein coverage of L_covered_/L = 90%, the neck radius decreases smoothly as the bending moduli ratio increases, approaching the spontaneous hemi-fission limit at a *k*_2_/*k* > 0.8 (pink line in Fig. 5A). (*ii*) For a protein coverage of 40% < L_covered_/L <70%, we observed a snap-through transition from a wide neck to a narrow neck below the hemi-fission threshold as the bending moduli ratio increased (green and blue lines in Fig. 5A). Based on our results, the snap-through transition shifts to larger *k*_2_/*k* ratio with decreasing the protein coverage (green and blue lines in Fig. 5A). (*iii*) For a protein coverage of L_covered_/L = 30%, our results showed that the radius of the neck slightly decreases as the bending moduli ratio increases up to *k*_2_/*k* = 4 (yellow line in Fig. 5A).

We also plotted the radius of the neck as a function of the bending moduli ratio for three different cubic lattice constants (Fig. 5B, L_covered_/L = 80%). We observed that for small cubic lattice constants (*a* = 10 and 15 nm), the constriction of mitochondrial membrane with increasing the anisotropic bending rigidity, is associated with a snap-through instability (blue and green lines in Fig. 5B). In contrast, for a large lattice constant (*a* = 20 nm), the radius of the neck slightly decreases as the bending rigidity ratio increases (pink line in Fig. 5B). The anisotropic bending rigidity of the membrane characterizes the magnitude of the membrane tension as we set 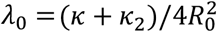. Previous studies have demonstrated that membrane tension plays an important role in governing the efficiency of mitochondrial fission ^12,60,72–75^. For example, in a low tension regime, complete fission occurs within a few minutes, while in a high membrane tension regime, it takes only a few seconds ^12,60,72–75^. In Fig. 5, the variation in *k*_2_/*k* from 0 to 4 corresponds to a membrane tension varying from *λ*_0_ ∼ 0.03 to *λ*_0_ ∼ 0.13 pN/nm. Morlot et. al., have experimentally shown that a high membrane tension, in the order of *λ*_0_ ∼ O (0.1) pN/nm, is required for successful mitochondrial fission within a few seconds ^74^. Thus, our results suggest that the dynamics of mitochondrial fission are governed by a complex interdependent relationship between membrane tension and the anisotropic bending rigidity of the membrane, covered by curvature-generating proteins.

#### Dynamin family of proteins can induce mitochondrial fission in a cardiolipin-dependent manner

Having established a mechanical framework to estimate the size of constricted necks by dynamin family of proteins from the cubic structures observed in SAXS experiments, it is interesting to compare the fission necks generated by Dnm1 and Drp1 in different lipid compositions. We have previously shown that Dnm1 can remodel lipid membranes to cubic structures with NGC ^21^. To test the membrane deformations induced by Drp1, we incubated Drp1 with small unilamellar vesicles (SUVs) at a protein-to-lipid (P/L) molar ratio of 1/1000 (Figs. 6A and S3). The SUVs were prepared using two different ternary phospholipid compositions of phosphatidylethanolamine (PE),phosphatidylcholine (PC), and cardiolipin (CL) at molar ratios of 75/15/10 and 75/5/20 to mimic the lipid compositions of mitochondrial membranes ^76^. We found that Drp1 restructured the lipid vesicles into a *Pn3m* cubic phase with a lattice constant of *a* = 19.075 nm and *a* = 34 nm for 75/5/20 and 75/15/10 PE/PC/CL, respectively (Figs. 6A and S3).

**Figure 6.**
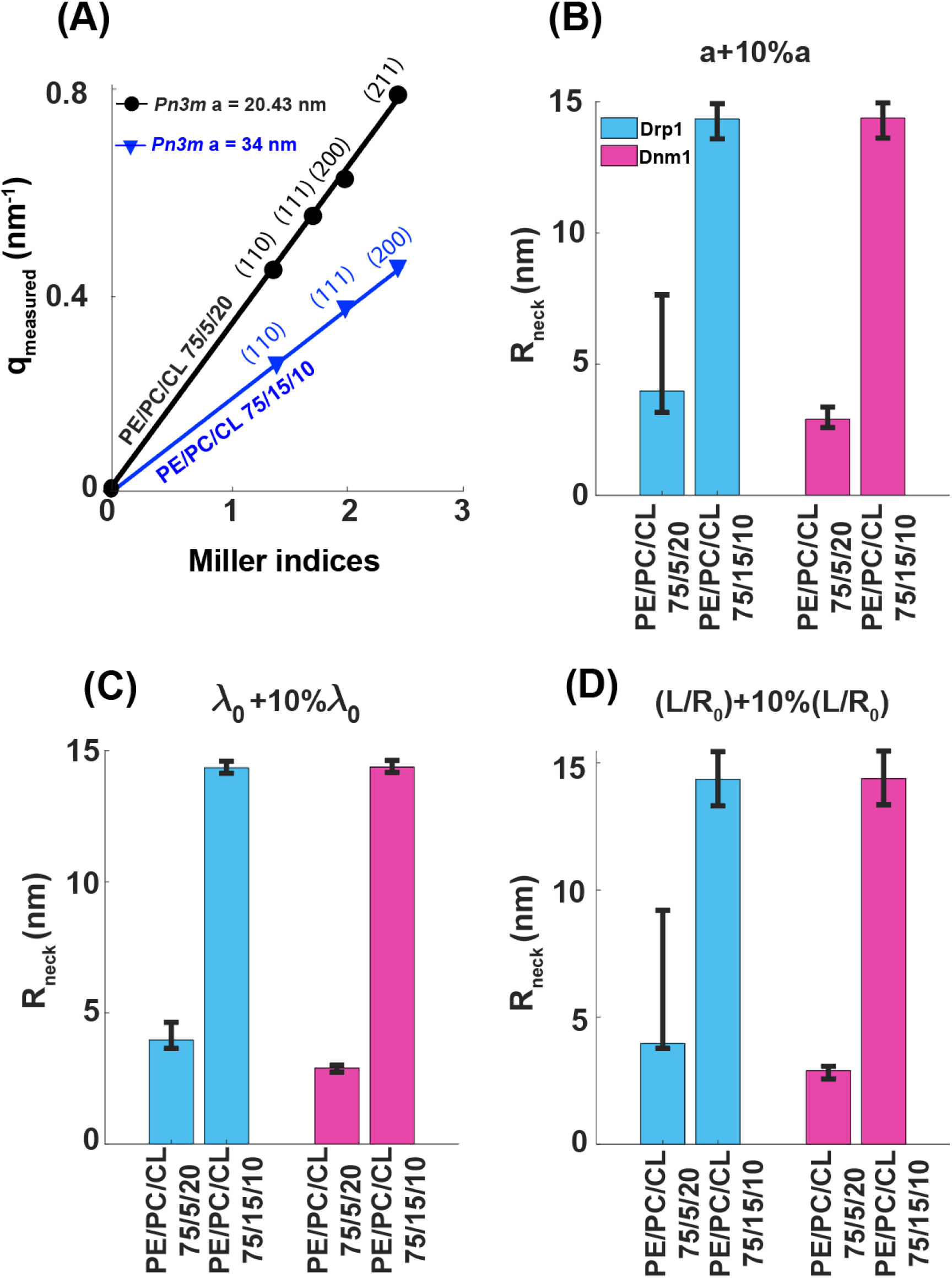
Variations in the size of scission necks induced by Drp1 and Dnm1 in two different lipid compositions. **(A)** Indexing of Drp1-induced cubic phases for 75/5/20 PE/PC/CL (black line) and 75/15/10 PE/PC/CL (blue line) model membranes and at a protein-to-lipid (P/L) molar ratio of 1/1000. Plots of the measured q positions (q_measured_) versus the assigned reflections in terms of Miller indices. Estimated radius of the constricted neck, accounting for 10% variations in **(B)** cubic lattice constants, **(C)** membrane tension (*λ*_0_ = 0.1 pN/nm), and **(D)** the length of the tubular membrane (L/R_0_ = 2).

We next applied our framework to explore the degree of neck constriction by Dnm1 and Drp1 in different lipid composition (Figs. 6B-6D). Using SAXS measurements as a baseline, we calculated the radius of constricted necks considering 10% variations in cubic lattice constants (representing technical variability), membrane tension (*λ*_0_ = 0.1 pN/nm), and the length of tubular membrane (L/R_0_ = 2) (Figs. 6B-6D). We assumed high membrane tension in our calculations, based on experimental observations that typical mitochondrial fission rate over a few seconds, is associated with a high membrane tension regime ^74^. We found that the efficiency of curvature-induced constriction highly depends on the composition of mitochondrial membrane (Fig. 6). Based on our results, in membranes with high cardiolipin concentration (PE/PC/CL 75/5/20), induced anisotropic curvature by Drp1 and Dnm1 can robustly constrict tubular membranes into narrow necks within the spontaneous hemi-fission range (Figs. 6B-6D). However, in membranes with low cardiolipin concentration, the induced curvature by Drp1 and Dnm1 slightly remodel membranes, forming wide necks with R_neck_ ∼ 15 nm (Figs. 6B-6D). Previous studies have shown that conical lipids such as cardiolipin play an important role in facilitating membrane-dynamin interactions and activation of the GTPase domain of fission proteins ^31–33,77^. Our estimated neck sizes induced by Drp1 and Dnm1 are ∼5x smaller than the diameters of constricted lipid tubules observed by electron microscopy for lipid-bound Drp1 oligomerization with 100% phospho-L-serine (PS) lipid composition ^78^. This could be due to the synergetic effects of lipid composition, mutually amplifying mechanical fission forces from membrane remodeling proteins, and Drp1 adaptors/recruiters that are responsible for bringing Drp1 to the specific sites of fission. For example, our results suggest that a modest reduction in the percentage of CL in lipid composition from 20% to 10% results in a nearly 5x increase in the radius of the scission neck induced by Drp1 and Dnm1 (Fig. 6), which can potentially make successful hemi-fission statistically improbable. Additionally, we observed that in membranes with high cardiolipin concentration, the radius of the constricted neck shows greater variability in response to changes in system properties, compared to compositions with lower cardiolipin concentration (Fig. 6). This can be an indication of the stochastic characteristic of the hemi-fission reaction, as suggested by both in *vivo* and in *vitro* experiments ^20,79–81^.

#### Coarse-grained molecular dynamics simulations show larger protein coverage induces stronger deviatoric curvature and its strength depends on membrane cardiolipin concentration

Our results from the continuum framework and SAXS data suggest that the dynamin family of proteins can constrict fission necks by imposing anisotropic curvature on the underlying tubular membrane. Based on our findings, the degree of anisotropic curvature generated by the dynamin family of proteins depends on the extent of membrane coverage by the protein (Fig. 2) and the percentage of cardiolipin in the lipid membrane (Fig. 6). To test this hypothesis, we conducted coarse-grained molecular dynamics simulations with the IDVD domains of Dnm1 on two different membrane lipid compositions: 75/5/20 PE/PC/CL and 75/15/10 PE/PC/CL (Fig. 7). In the first simulations, we placed two IDVD domains of Dnm1 on a 75/15/10 PE/PC/CL membrane in such a way that the variable loops are closer to the membrane surface (Fig. 7A). The system was equilibrated for 2 μs, and the equilibrated trajectory was used to compute the deviatoric curvature generated on the membrane. Details of the simulation protocol and calculations of deviatoric curvature are described in the Supplementary Material. Fig. 7B shows that the IDVD domain of Dnm1 can generate deviatoric curvature on the membrane surface. We then increased the protein coverage on the membrane (corresponding to the L_covered_ of the theoretical model) by placing four IDVD domains on the membrane surface and observed ∼19% increase in the magnitude of induced deviatoric curvature on the membrane surface (Fig. 7C). This is consistent with the prediction of our theoretical model that larger protein coverage induces narrower mitochondrial fission necks (Fig. 2A).

**Figure 7.**
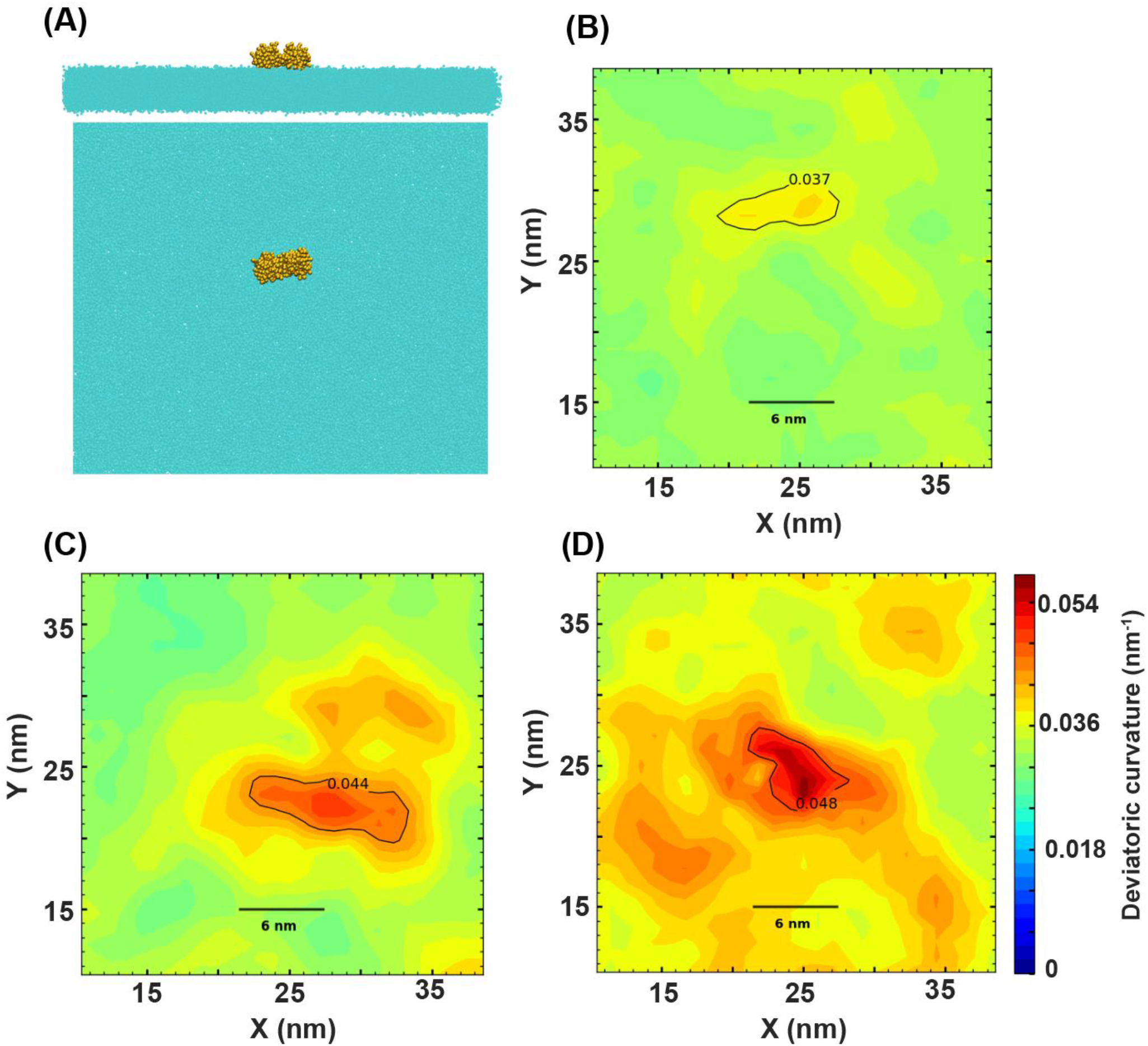
Comparison of the magnitude of deviatoric curvature induced by dynamin with changing the protein coverage and the cardiolipin concentration of lipid membrane using molecular dynamics simulations. **(A)** The snapshot shows the top and side view of the lipid membrane containing two IDVD domains. Deviatoric curvature induced by **(B)** two and **(C)** four IDVD domains on a 75/15/10 PE/PC/CL lipid membrane. **(D)** Deviatoric curvature induced by two IDVD domains on a 75/5/20 PE/PC/CL membrane.

Next, we investigated the effect of lipid composition, particularly cardiolipin, on the degree of deviatoric curvature generated by Dnm1. We placed two IDVD domains on the surface of a 75/5/20 PE/PC/CL lipid membrane (Fig. 7D). Our results show that the deviatoric curvature induced by Dnm1 on this lipid membrane is significantly (∼30%) higher than the deviatoric curvature generated on a 75/15/10 PE/PC/CL lipid membrane (Figs. 7B and 7D). This is also in agreement with our theoretical predictions and SAXS experiments, which show that a modest increase in the percentage of cardiolipin in the lipid composition results in a significant decrease in the radius of the scission neck induced by Dnm1 proteins. Additionally, we observed that the induced deviatoric curvature on the 75/5/20 PE/PC/CL lipid membrane extends to a relatively larger distance compared to that on the 75/15/10 PE/PC/CL lipid membrane (Figs. 7B and 7D), which might be due to increased membrane fluidity with increasing cardiolipin concentration ^82^. A previous study has suggested that the mechanical stability of lipid membranes decreases with increasing cardiolipin concentration ^82^. This implies that a higher concentration of cardiolipin in the mitochondrial membrane can facilitate the process of fission neck constriction, as predicted by our coarse-grained simulations and theoretical model.

## Discussion

Mitochondria are dynamic organelles that constantly undergo fission and fusion processes. The balance between these two processes plays an important role in modulating the shape, distribution of mitochondria, and bioenergetics ^1–3^. For example, excessive fission of mitochondria has been implicated in neurodegenerative diseases ^4–6^. The dynamin family of proteins (Dnm1 in yeast and its conserved human homologues, Drp1) is key molecular machinery responsible for regulating mitochondrial fission ^1,7–10^. Despite three decades of experimental work, the mechanical principle underlying membrane scission is still unclear ^12^. Dynamin is a mechano-chemical enzyme, and current models for dynamin-driven fission can broadly be categorized into two main classes: (*I*) the two-step model and (*II*) the constrictase model ^12^. In the two-step model, dynamin’s helical assembly first slowly constricts the mitochondrial membrane into a tubular structure, followed by a rapid transition to scission, driven by GTP hydrolysis-induced oligomer disassembly ^12,17–19^. However, this process requires high cooperativity among dimers, which is inconsistent with the low measured Hill coefficient, and the suggested disassembly could cause tubule widening due to the slower rate of dynamin disassembly compared to membrane viscoelasticity. On the other hand, in the constrictase model, dynamin serves as a molecular motor protein where GTP hydrolysis prompts the sliding of helices, leading to spontaneous hemi-fission. However, in this model, how disassembly occurs and how such disassembly contributes to scission remain a matter of debate. In particular, the required number of interacting helicoidal coils to generate the force necessary for scission seems to be variable, exhibiting a degree of stochasticity. These open challenges in each dynamin fission model are further complicated by the extreme changes in the geometry of the mitochondrial membrane during the fission process, transitioning from a tubular structure to a catenoid-shaped neck with a large NGC.

Our results from the mechanical framework combined with SAXS measurements suggest that the fission protein machinery of dynamin governs mitochondrial fission through both its helix-driven mechanical confinement of the tubular membrane and its synergistic curvature-driven snap-through instability. A previous study by Irajizad et al. suggested that actin-mediated forces, combined with the curvature induced by conical lipids localized in the pinching domain, can trigger buckling instability during mitochondrial fission ^25^. Here, we propose that the NGC induced by dynamin alone is sufficient to drive the snap-through instability and generate super-constricted fission necks below the membrane’s hemi-fission threshold. It is interesting to note that the strongest NGC-inducing domain in Dnm1 is not the variable domain facing the confined membrane tubule, but rather the GTPase domain more distal from it ^21^. In support of this, we showed in the companion paper that the hydrolysis-driven oligomer disassembly and concomitant creation of a population of mobile dynamins can significantly amplify NGC generation by presentation of the free NGC-generating GTPase domain in the vicinity of the mitochondrion.

Previous studies have proposed the presence of mechanical instability during dynamin constriction due to an increase in the helical pitch of the dynamin assembly or an adjustment in the spontaneous curvature of the neck membrane to favor negative values ^83,84^. Our results introduce a new perspective; the collar pressure induced by helical assemblies of dynamin smoothly constricts the mitochondrial membrane into narrow necks (Fig. 3), while curvature-driven remodeling is associated with a snap-through transition (Fig. 2). This difference resembles the observed dynamics of slow and smooth membrane neck constriction facilitated by dynamin assembly, followed by the fast mitochondrial fission step after GTP hydrolysis, which induces helix disassembly-a phenomenon broadly observed experimentally. These effects essentially recapitulate major observed features of the two-step model of dynamin-induced mitochondrial fission. It should be noted that our focus here was on the equilibrium shapes of the membrane, and we did not model the exact dynamical transitions between stages. However, from a mechanical perspective, we can expect that the curvature-induced “snap-through” transition to occur within the membrane relaxation timescale (∼ms) ^85,86^, which is comparable to experimental observations for the fast induced fission by dynamin disassembly in the two-step model ^12^. Importantly, we find that longer tubular membranes (Length/Radius > 1.8) are more prone to undergoing the fission process in response to the NGC generated by the dynamin family of proteins (Fig. 4). Our results demonstrate that, under a large induced NGC, a snap-through transition occurs in the hemi-fission region as the length of a tubular membrane increases (Fig. 4). This length-associated snap-through instability may serve as the driving mechanism for spontaneous hemi-fission in the constrictase model, considering that the tubular membrane elongates with constriction ^63^. What’s more, our results here suggest that there is an intrinsic stochasticity to the fission process, since in principle different combinations of dynamin helical oligomer confinement, free dynamin mediated membrane remodeling, and membrane tubule length can lead to completion of fission.

In current models, the precise number of dynamin helical turns required to induce mitochondrial fission is not so clear ^12^. Based on our results, the dynamin helical scaffolds only need to constrict the tubular membrane into a wide neck with a radius of R_neck_ ∼ 8 nm, at which point the curvature-driven instability can complete the scission (Figs. 2 and 3). According to our calculations, a collar pressure of ∼ 4 pN/nm is required to constrict the tubular membrane to the radius of R_neck_ ∼ 8 nm (Fig. 3). Considering each helix turn of dynamin can exert a compressive force of ∼ 110 pN ^63^, and the helical pitch varies between 10-20 nm ^14^, our results suggest that 1 to 3 dynamin helical turns would be sufficient to drive mitochondrial fission. This is in accord with *in vivo* studies that have shown that even a single helical turn is sometimes sufficient for efficient fission ^81,87^.

The results presented here comprise an important step toward deciphering the intricate mechanochemical principles underlying mitochondrial fission. Our findings can be a motivation for future important studies to develop quantitative relationships between GTP-driven disassembly of dynamin and generation of membrane-destabilizing curvature. Particularly, in the companion paper, we elucidate how dynamin mediates fission across a heterogeneously uncorrelated population in response to varying levels of GTP hydrolysis. This could provide valuable insights for understanding the relationship between GTP-driven disassembly and scission, the precise role of the energy induced by GTP hydrolysis in enabling dynamin-driven fission, and ultimately comparing endocytic dynamin versus mitochondrial dynamin.

## Acknowledgments

This work was supported by American Heart Association AHA966662 grant and NSF DMR2325840 (G.C.L.W), National Institutes of Health grant R01GM067180 (RBH, GCLW), and Vascular Biology Training Grant T32 HL069766-21 (H. A). T.M gratefully acknowledges the support from the Government of India: Science and Engineering Research Board via Sanction No. SRG/2022/000548. We thank the Stanford Synchrotron Radiation Lightsource (SSRL) (Menlo Park, CA, USA) for access to beamline 4-2. Use of the SSRL, SLAC National Accelerator Laboratory, is supported by the U.S. Department of Energy, Office of Science, Office of Basic Energy Sciences under contract no. DE-AC02-76SF00515. The SSRL Structural Molecular Biology Program is supported by the U.S. Department of Energy, Office of Biological and Environmental Research, and by the National Institutes of Health, National Institute of General Medical Sciences (including P30GM133894). T.M is grateful for the computational resources provided by PARAM Sanganak under the National Supercomputing Mission, Government of India, at the Indian Institute of Technology, Kanpur.

