## Supplementary material for "Dynamins combine mechano-constriction and membrane remodeling to enable two-step mitochondrial fission via a ‘snap-through’ instability"

### Membrane mechanics

We modeled the lipid bilayer as a continuous elastic shell with negligible thickness [1,2]. Considering the stretching modulus of the lipid bilayer is an order of magnitude larger than the membrane bending modulus [35], we assumed that the membrane is locally inextensible. Assuming the system is in mechanical equilibrium at all times, the total energy of the system including the elastic storage energy of the membrane ($E_{elastic}$) and the work done by the applied forces by dynamin helical rings ($E_{force}$) is given by [3–5]

$$E = E_{elastic} - E_{force}, \tag{S1}$$

where

$$E_{elastic} = \int_{\omega} W(H, D; \theta^{\alpha}) + \lambda(\theta^{\alpha}) da - pV \quad \text{and} \tag{S2a}$$

$$E_{force} = \int_{\omega} \boldsymbol{\tau}(\theta^{\alpha})(\boldsymbol{r} - \boldsymbol{r_0})\, da, \tag{S2b}$$

where $\omega$ is the total membrane surface area, $W$ is the energy density, $\theta^{\alpha}$ represents the surface coordinate where $\alpha \epsilon \{1,2\}$, $H$ is the mean curvature of the surface, and $D$ is the curvature deviator. $\lambda$ is the surface tension field which is a Lagrange multiplier associated with the local area constraint, $p$ is the transmembrane pressure, and $V$ is the enclosed volume. $\boldsymbol{\tau}$ is the compressive force density on the membrane induced by helical arrangements of dynamin proteins, $\boldsymbol{r}$ is the position vector in the current configuration, and $\boldsymbol{r_0}$ is the position vector in the reference frame.

A balance of forces normal to the membrane yields the "shape equation" given as [6,7]

$$\underbrace{\tfrac{1}{2}\left[W_D\left(\xi^{\alpha}\xi^{\beta} - \mu^{\alpha}\mu^{\beta}\right)\right]_{;\beta\alpha} + \tfrac{1}{2}W_D\left(\xi^{\alpha}\xi^{\beta} - \mu^{\alpha}\mu^{\beta}\right)b_{\alpha\gamma}b_{\beta}^{\gamma} + \Delta\left(\tfrac{1}{2}W_H\right) + W_H(2H^2 - K) - 2HW}_{\text{Elastic effects}} =$$

$$\underbrace{p + 2H\lambda}_{\substack{\text{Capillary}\\ \text{effects}}} + \underbrace{\boldsymbol{\tau}.\boldsymbol{n}}_{\substack{\text{Compressive stress}\\ \text{induced by Dnm1}}} \tag{S3}$$

where $b_{\alpha\gamma}$ is the coefficients of the second fundamental form, $b_{\beta}^{\gamma}$ is the mixed components of the curvature, and $(.)_{;\alpha}$ is the covariant derivative. $W_D$ and $W_H$ are the partial derivatives of the energy density $W$, $K$ is the Gaussian curvature, and $\Delta(.)$ is the surface Laplacian. $\xi^{\alpha}$ and $\mu^{\alpha}$ are the projections of $\boldsymbol{\xi}$ and $\boldsymbol{\mu}$ along the tangent vector on the surface with

$$\xi^{\alpha} = \boldsymbol{\xi}.\boldsymbol{a}^{\boldsymbol{\alpha}}, \tag{S4a}$$

$$\mu^{\alpha} = \boldsymbol{\mu}.\boldsymbol{a}^{\boldsymbol{\alpha}}, \tag{S4b}$$

where $\boldsymbol{a}^{\boldsymbol{\alpha}}$ is the contravariant basis vectors, $\boldsymbol{\xi}$ is a unit vector representing the orientation of a one-dimensional curve on the surface, which is tangential to the protein coat, and $\boldsymbol{\mu}$ is a unit vector defined as

$$\boldsymbol{\mu} = \boldsymbol{n} \times \boldsymbol{\xi}, \tag{S5}$$

where $\boldsymbol{n}$ is the normal vector to the surface.

A balance of forces tangent to the membrane yields the spatial variation of membrane tension given as [6,7]

$$\lambda_{,\alpha} = -W_{,\alpha|exp} - \boldsymbol{\tau}.\boldsymbol{a_s}, \tag{S6}$$

where $(.)_{\alpha}$ is the partial derivative, $(.)_{|exp}$ denotes the explicit derivative with respect to coordinate, and $\boldsymbol{a_s}$ is a tangent vector on the surface.

We modeled the energy density of the membrane-dynamin interactions using the modified version of Helfrich energy including the deviatoric curvature given as [5,8–11]

$$W = \kappa(H - C_0)^2 + \kappa_2(D - D_0)^2 \tag{S7}$$

where $C_0$ and $D_0$ are the induced spontaneous isotropic and deviatoric curvatures by dynamin-related GTPase proteins, respectively. $\kappa$ is the membrane bending rigidity associated with isotropic curvature and $\kappa_2$ is the membrane bending rigidity associated with anisotropic curvature. Assuming the induced isotropic curvature is negligible compared to the deviatoric curvature ($C_0 \ll D_0$), we can substitute Eq. S7 in Eqs. S3 and S6 and rewrite the forces balances as

$$\kappa_2\left[(D - D_0)\left(\xi^{\alpha}\xi^{\beta} - \mu^{\alpha}\mu^{\beta}\right)\right]_{;\beta\alpha} + \kappa_2(D - D_0)\left(\xi^{\alpha}\xi^{\beta} - \mu^{\alpha}\mu^{\beta}\right)b_{\alpha\gamma}b_{\beta}^{\gamma} + \kappa\Delta(H - C_0) + 2\kappa(H - C_0)(2H^2 - K) - 2HW = p + 2H\lambda + \boldsymbol{\tau}.\boldsymbol{n}\,, \tag{S8}$$

and

$$\lambda_{,\alpha} = 2\kappa_2(D - D_0)\frac{\partial D_0}{\partial \theta^{\alpha}} - \boldsymbol{\tau}.\boldsymbol{a_s}. \tag{S9}$$

**Parametrization in axisymmetric coordinates**

For ease of computation, we assumed that the tubular membrane and the constricted necks are rotationally symmetric. This allows us to parametrize the normal and tangent vectors on the surface given as

$$\boldsymbol{n} = -\sin(\psi)\boldsymbol{e_r} + \cos(\psi)\boldsymbol{k} \quad \text{and} \quad \boldsymbol{a_s} = \cos(\psi)\boldsymbol{e_r} + \sin(\psi)\boldsymbol{k}, \tag{S10}$$

where $\boldsymbol{e_r}$ and $\boldsymbol{k}$ are the coordinate basis, and $\psi$ is defined as an angle made by the tangent with respect to the horizontal. We can now write the radial distance from the axis of rotation ($R(s)$), and the elevation from the reference plane ($Z(s)$) as a function of one parameter arclength ($s$) given as

$$R'(s) = \cos(\psi),$$

$$Z'(s) = \sin(\psi), \tag{S11}$$

where $(.)' = \frac{d(.)}{ds}$. Defining $M = \frac{R}{2}[(W_H)' - (W_D)'] - \cos(\psi)W_D$, we can simplify the normal and tangential force balances (Eqs. S8 and S9) to the system of first order differential equation given as

$$R'(s) = \cos(\psi),\ \ Z'(s) = \sin(\psi),\ \ R\psi' = 2RH - \sin(\psi),$$

$$R(\kappa + \kappa_2)H' = M - R\kappa_2 D_0' - 2\kappa_2\cos(\psi)\, D_0,$$

$$\frac{M'}{R} = p + 2\kappa H\left[\frac{\kappa_2}{\kappa}D_0^2 + \frac{\lambda}{\kappa} - \left(1 + \frac{\kappa_2}{\kappa}\right)\left(H - \frac{\sin(\psi)}{R}\right)^2\right] + \tau\sin(\psi),$$

$$\lambda' = 2\kappa_2\left(\frac{\sin(\psi)}{R} - H - D_0\right)D_0' - \tau\cos(\psi). \tag{S12}$$

In order to solve the system of partial differential equations in Eq. S12, we prescribe six boundary conditions as follow:

$$Z(0) = 0,\ \psi(0) = \frac{\pi}{2},\ M(0) = 0,\ R(s_{max}) = R_0,\ \ \psi(s_{max}) = \frac{\pi}{2},\ \lambda(s_{max}) = \lambda_0, \tag{S13}$$

where $s_{max}$ is the maximum length of the computational domain and $\lambda_0$ is the prescribed membrane tension at the boundary.

In our numerical calculations, to have a sharp but smooth transition at the boundaries of covered area by the curvature-generating proteins, we prescribed the induced anisotropic curvature using a hyperbolic tangent function given as

$$D_0 = \frac{D_m}{2}[\tanh(g(s - s_0))], \tag{S14}$$

where $g$ is a constant and $s_0$ represents the covered length of the tube by proteins.

**Nondimensionalization**

We nondimensionalized the system of equations (Eq. S12) using two positive constants, the radius of the membrane tubule ($R_0$) and the membrane bending rigidity ($\kappa$), and defining the following dimensionless variables given as

$$t \equiv \frac{s}{R_0},\ r \equiv \frac{R}{R_0},\ z \equiv \frac{Z}{R_0},\ h \equiv HR_0,\ d_0 \equiv D_0R_0,\ m = \frac{MR_0}{\kappa},\ \tilde{p} = \frac{pR_0^3}{\kappa},\ \tilde{\lambda} \equiv \frac{\lambda R_0^2}{\kappa},\ \tilde{\kappa} \equiv \frac{\kappa_2}{\kappa},\ \tilde{\boldsymbol{\tau}} \equiv \frac{\boldsymbol{\tau} R_0^3}{\kappa}. \tag{S15}$$

Rewriting Eqs. S12 and S13 in terms of the dimensionless variables, we get

$$\dot{r} = \cos(\psi),\ \ \dot{z} = \sin(\psi),\ r\dot{\psi} = 2rh - \sin(\psi),$$

$$r(1+\tilde{\kappa})\dot{h} = m - r\tilde{\kappa}D_0' - 2\tilde{\kappa}\cos(\psi)\, d_0,$$

$$\frac{\dot{m}}{r} = \tilde{p} + 2h\left[\tilde{\kappa}d_0^2 + \tilde{\lambda} - (1+\tilde{\kappa})\left(h - \frac{\sin(\psi)}{r}\right)^2\right] + \tilde{\tau}\sin(\psi),$$

$$\dot{\lambda} = 2\tilde{\kappa}\left(\frac{\sin(\psi)}{r} - h - d_0\right)\dot{d}_0 - \tilde{\tau}\cos(\psi), \tag{S16}$$

where $(\dot{X}) = \frac{d(X)}{dt}$ and the boundary conditions simplify as

$$z(0) = 0,\ \psi(0) = \frac{\pi}{2},\ m(0) = 0,\ r(t_{max}) = 1,\ \psi(t_{max}) = \frac{\pi}{2},\ \tilde{\lambda}(t_{max}) = \tilde{\lambda}_0. \tag{S17}$$

**Radius of a cylinder from energy minimization**

Let us consider a tubular membrane with a radius $R_0$ and a height of 2L under tension $\lambda_{cylinder}$. By using the Helfrich energy in Eq. S7 and assuming no spontaneous curvatures ($C_0 = 0$ and $D_0 = 0$), we can simplify the total energy of the membrane as

$$E_{cylinder} = \left(\frac{\kappa+\kappa_2}{4R_0^2} + \lambda_{cylinder}\right)4\pi R_0 L. \tag{S18}$$

By solving $\partial E_{cylinder}/\partial R_0$ = 0 for mechanical equilibrium, we can find the relationship between the radius of the tubule, membrane bending rigidity, and tension, as given by [5,12]

$$R_0 = \frac{1}{2}\sqrt{\frac{\kappa+\kappa_2}{\lambda_{cylinder}}}. \tag{S19}$$

For a tubular membrane under a uniform radial force density, $\tau$, along its length, Eq. S18 is modified as follows [5]

$$E_{cylinder} = \left(\frac{\kappa+\kappa_2}{4R_0^2} + \lambda_{cylinder} + \tau\right) 4\pi R_0 L, \tag{S20}$$

and the radius of cylinder in mechanical equilibrium is given by [5]

$$R_0 = \frac{1}{2}\sqrt{\frac{\kappa+\kappa_2}{\lambda_{cylinder}+\tau}}\,. \tag{S21}$$

**Coarse-grained Molecular Dynamics simulations**

All coarse-grained molecular dynamics simulations were performed using the MARTINI-2.2 forcefield [13]. Initial structures of the model membranes were constructed using the "Martini Builder" module of the CHARMM-GUI software [14,15]. The model membranes were composed of dioleyl phosphoethanolamine (DOPE), dipalmitoyl phosphocholine (DOPC), and cardiolipin (CL) lipids with a molar ratio of 75/15/10 and 75/5/20, respectively. Initial coordinates of the Pleckstrin Homology (PH) domain were taken from the protein data bank (PDB ID: 1dyn). We first placed a protein dimer on top of a 75/15/10 DOPE/DOPC/CL membrane which was solvated using MARTINI coarse-grained water along with 10% anti-freezing beads. Appropriate numbers of $Na^+$and $Cl^-$ ion beads were then added to maintain 150 mM salt concentration. The solvated protein-membrane complex structure was then energy minimized using the steepest descent and conjugate gradient methods to remove any bad contacts between the solute and solvate beads. The energy minimized structure was then subject to short equilibration using constant-volume-constant-temperature (NVT) and constant-pressure-constant-temperature (NPT) ensembles to achieve correct density. Subsequently, 2 µs long production run was performed at 303 K temperature and atmospheric pressure. A velocity-rescale thermostat [16] with a time constant of 1ps, was used to maintain the temperature of the system and the pressure was maintained using the Parrinello-Rahman barostat [17] with a time constant of 12 ps. The simulation time step was set to 5 fs for the numerical integrations, and the system coordinates were saved at every 100 ps. The tertiary structures of the proteins were maintained using an elastic network (ELNeDyn2.2) [18] with a force constant of 500 $kJ.mol^{-1}.nm^{-2}$ during all the simulations. All simulations were performed using GROMACS software [19]

We employed MDAnalysis tool [20] and the VTK library [21] of Python to compute the deviatoric curvature induced by the proteins. We first obtained a triangulated surface using the Delaunay Triangulation of the "PO4" beads from the upper leaflet of the bilayer. This surface was then smoothed using the Laplacian smoothing algorithm with a relaxation factor of 0.5 and 10 iterations for each frame. The mean (H) and Gaussian (K) curvatures were calculated at each vertex point using the VTK library functions. The deviatoric curvature (D) was then calculated from the mean and Gaussian curvatures using the relationship $D = [|H^2 - K|]^{1/2} = \frac{1}{2}[|C_1 - C_2|]$, where $C_1$ and $C_2$ are the principal curvatures. To investigate the effect of protein coverage on the deviatoric curvature, four variable domains from the dynamin protein quaternary structure (PDB ID: 4uud) were placed on the 75/15/10 DOPE/DOPC/CL membrane. Finally, we considered a 75/5/20 DOPE/DOPC/CL membrane to investigate the effect of cardiolipin on the deviatoric curvature induced by the proteins. All the protein membrane complex structures were equilibrated using the above-mentioned simulation protocol.

**Supplementary figures**

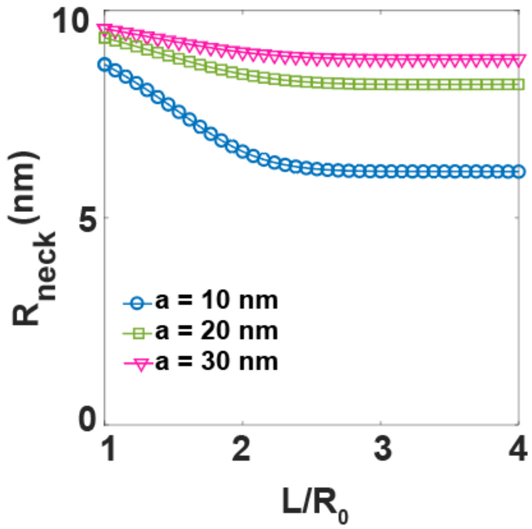


**Figure S1.** Decrease in the radius of mitochondrial constricted neck as the function L/R0. Here, we fixed $L_{covered}$ = 36 nm for all tubule heights.

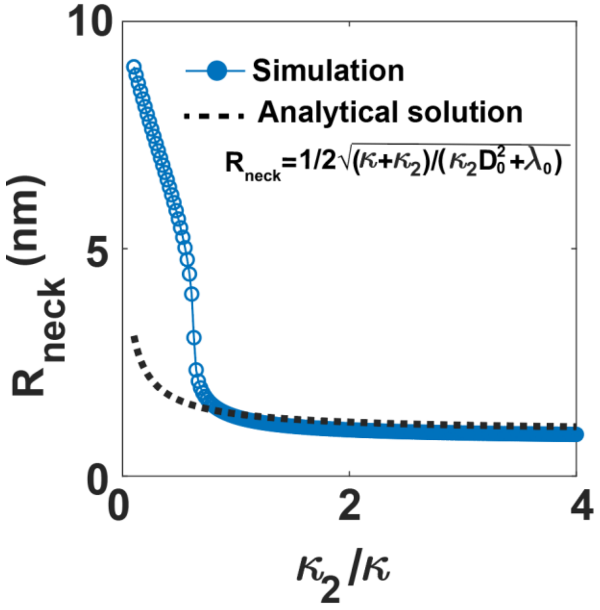


**Figure S2.** Decrease in the radius of mitochondrial constricted neck as the function anisotropic to isotropic bending rigidity ratio ($\kappa_2/\kappa$). The dotted black line represents the analytical expression for the equilibrium radius of a tubular membrane in the presence of deviatoric curvature which is given as $R_{tube}$ = $1/2\sqrt{(\kappa+\kappa_2)/(\kappa_2 D_0^2+\lambda_0)}$. The blue line indicates the results from numerical simulations ($a$ = 10 nm and $L_{covered}$ = 90%). Based on our results, for large $\frac{\kappa_2}{\kappa} > 1$, there is a good agreement between the analytical solution and numerical results. However, for smaller $\frac{\kappa_2}{\kappa} < 1$, the analytical solution underestimates the radius of the neck.

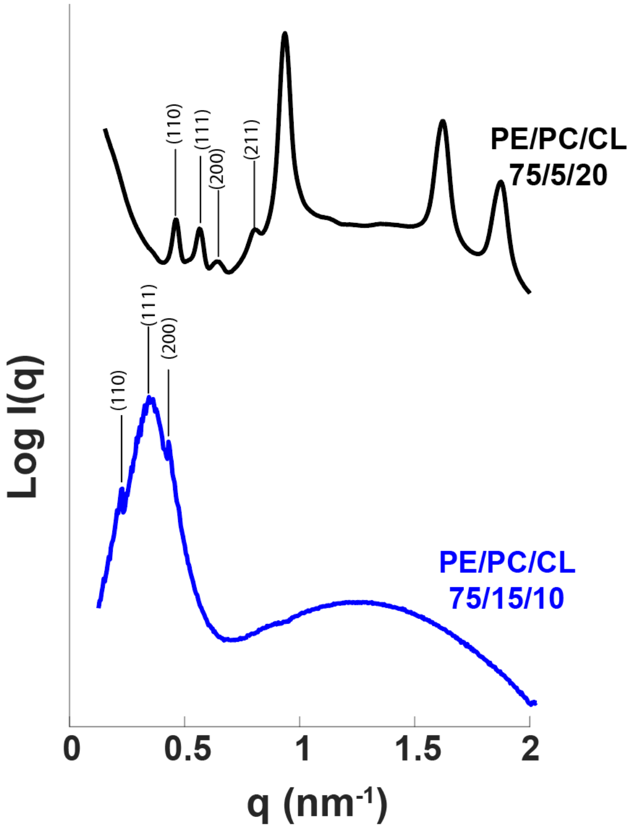


**Figure S3.** SAXS spectra from 75/15/10 and 75/5/20 PE/PC/CL model mitochondrial membranes incubated with Drp1 at a protein-to-lipid (P/L) molar ratio of 1/1000. The peaks correspond to a *Pn3m* cubic phase with a lattice constant of $a$ = 20.43 nm and $a$ = 34 nm for a lipid composition of 75/5/20 and 75/15/10 PE/PC/CL, respectively.

21. vtk: VTK is an open-source toolkit for 3D computer graphics, image processing, and visualization.
